# Multi-spectral optoacoustic microscopy driven by gas-filled hollow-core fiber laser pulses

**DOI:** 10.1101/2025.05.27.656278

**Authors:** Cuiling Zhang, Marcello Meneghetti, J. E. Antonio-Lopez, Rodrigo Amezcua-Correa, Yazhou Wang, Christos Markos

**Affiliations:** DTU Electro, Technical Universisty of Denmark, DK-2800 Kgs. Lyngby, Denmark; Department of Neuroscience, University of Copenhagen, Blegdamsvej 3B, DK-2200 Kbh N, Copenhagen, Denmark; CREOL, The College of Optics and Photonics, University of Central Florida, Orlando, Florida 32816, USA; NORBLIS ApS, Virumgade 35D, DK-2830 Virum, Denmark

## Abstract

Multi-spectral optoacoustic microscopy (MS-OAM) requires high-performance light sources capable of delivering multiple intense spectral lines precisely matched to the absorption characteristics of selected biomolecules. We present a gas-filled anti-resonant hollow-core fiber (ARHCF) laser source optimized for near-infrared (NIR) MS-OAM. The hydrogen (H_2_)-filled ARHCF laser emits multiple spectral lines with high pulse energy and narrow linewidths (<0.1 nm) across a broad spectral range (∼1100 nm to ∼2200 nm). Several Raman laser lines were generated to overlap with key biomolecular absorption bands, including lipids (1210 nm and 1700 nm), collagen (∼1540 nm), and water (∼1400 nm and ∼1900 nm). We demonstrate the system’s performance by mapping absorbers in the first and second overtone regions of hair, pig tissue, and collagen samples. This work aims to bring the gas-filled fiber technology in MS-OAM applications and paves the way for high-resolution, label-free bio-imaging across extended infrared and ultraviolet regimes.

## 1. Introduction

Optoacoustic (also known as photoacoustic) microscopy (OAM) is a hybrid, non-invasive imaging technique that offers deep tissue penetration by detecting ultrasound waves generated through wavelength-dependent chromophore absorption [1,2]. In recent years, near-infrared (NIR) OAM operating in the first and second overtone regions has gained significant attention within the optoacoustic (OA) research community [3–6], as it targets the vibrational signatures of key biomolecules such as water (∼1400 nm and ∼1900 nm) [7,8], proteins (∼1500 nm, corresponding to combination overtones of the Amide I and II bands), and lipids (1200 nm and 1700 nm) [9]. Owing to this abundance of distinct absorption features, there is growing interest in employing multiple wavelengths to extract detailed spectral and functional information in OAM [5,6].

Achieving effective multi-spectral OA imaging requires laser sources that deliver multiple excitation wavelengths across the desired spectral range with enough pulse energy. Conventional OAM systems often rely on multiple independent laser sources to meet these needs, increasing system complexity and cost [10–13]. While rare-earth-doped fiber lasers offer multiwavelength laser emission, their tuning range is limited to tens of nanometers due to the narrow gain bandwidth of the dopants [14,15]. More recently, nanosecond multiwavelength pulsed lasers based on stimulated Raman scattering (SRS) in solid-core fibers have been developed for OA imaging [9,16–19] (Supplementary Note 1). However, these systems are constrained by the small Raman shift coefficient of silica, limiting their wavelength coverage to ∼113 nm (13.2 THz), and by their broad linewidths resulting from wide Raman gain bandwidths [9]. Supercontinuum (SC) light sources have also been explored for MS-OAM [17,20–23], as they can span from the ultraviolet (UV) to the mid-infrared (MIR), covering most biomolecular absorption bands. However, the low pulse energy density (PED) of SC sources requires the use of broad bandpass filters (∼tens of nanometers) to isolate sufficient energy to excite strong acoustic signals, which compromises spectral resolution [20].

Gas-filled anti-resonant hollow-core fiber (ARHCF) Raman lasers represent a promising alternative for generating multiwavelength laser sources [24–30]. These fibers support broad transmission from UV to MIR due to their hollow-core structure, which minimizes propagation loss [29]. Consequently, they enable generation of cascaded Raman and anti-Stokes lines over a wide spectral range [27–31]. Moreover, using gas as the Raman medium offers two main advantages: (i) narrow Raman gain bandwidth for producing narrow-linewidth laser pulses, and (ii) a large Raman shift coefficient, enabling wide spectral tunability from the pump wavelength. These features position ARHCF technology as a compelling solution for MS-OAM. For instance, recent demonstrations include gas-filled Raman lasers emitting multiple narrow lines from 1.5 to 2.4 µm [29].

Despite this potential, multi-wavelength gas-filled ARHCF lasers have yet to be explored in optoacoustic imaging. In this work, we report OAM using a hydrogen (H_2_)-filled ARHCF laser source that covers a broad wavelength range overlapping with the first and second overtone regions. We demonstrate that this source delivers high-energy, narrow-linewidth (<0.1 nm) nanosecond pulses suitable for high-resolution OAM. Furthermore, we implement the laser in a scanning transmission-mode OAM system and employ it for label-free imaging of lipid, melanin, and collagen distributions.

## 2. Materials and methods

### 2.1 Gas-filled ARHCF laser source

Figure 1 shows the schematic of the ARHCF pulsed laser source used in this study. The system employs a custom-made all-fiber Yb-doped fiber amplifier that emits pulse trains at two wavelengths, 1060 nm and 1044 nm, with a repetition rate of 1 kHz and pulse duration of 3.7 ns [31]. The corresponding pulse energies were measured to be approximately 98 µJ and 75 µJ, resulting to peak powers of ∼26.5 kW and ∼20.3 kW, respectively, as measured with a pyroelectric energy meter (PE9-ES-C, Ophir Optronics). The pump laser exhibits a narrow spectral linewidth of < 0.2 nm at both wavelengths, which facilitates the generation of narrow-linewidth Raman pulses in this work.

**Fig. 1.**
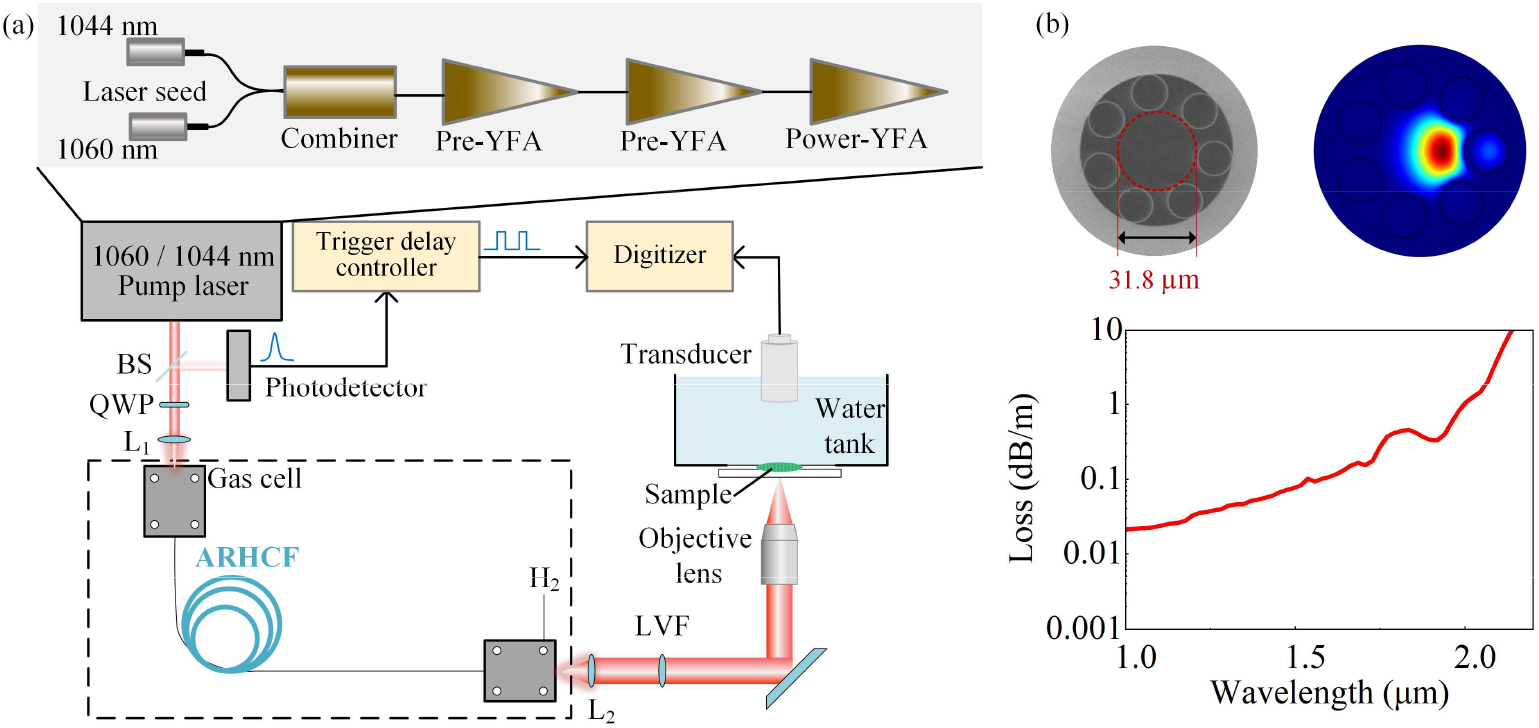
(a) Experimental setup. BS: beamsplitter; QWP: quarter-wave plate; ARHCF: anti-resonant hollow-core fiber; L_1_ and L_2_: focusing lenses; LVF: linear variable filter. YFA: Yb-doped fiber amplifier. (b) Left: SEM image of ARHCF. Right: The simulated fundamental mode field distribution of the ARHCF at 1.5 µm with a fiber bending radius of 15 cm. Bottom: The simulated fiber bend losses of the fundamental mode as a function of wavelength with a fiber bending radius of 15 cm [31].

The central wavelengths of the pump laser can be tuned over a ∼1 nm range by thermally adjusting the seed laser temperatures between 20 °C and 35 °C, allowing wavelength-tunable Raman output. The pump beams are launched into a 16-meter-long, H_2_-filled, 7-tube nodeless ARHCF with core diameter of 32.8 µm, capillary diameter of 16.1 µm, and wall thickness of ∼323 nm (Fig. 1(b)). This fiber structure supports low-loss transmission of fundamental mode in the NIR (1 µm to ∼1.6 µm), with losses below 0.1 dB/m [31], as shown in Fig. 1(b). Losses increase towards longer wavelengths, limiting the output to approximately 2.2 µm in this configuration. The ARHCF is sealed within two high-pressure custom-made gas cells, where light is coupled in and out through optical windows and focusing lenses [28]. The coupling efficiency from the pump laser into the ARHCF is approximately 62%. A qarter-wave plate (QWP) is placed before the input window to adjust the pump pulse polarization for optimal Raman generation.

### 2.2 Transmission-mode NIR-OAM

The generated beams by the ARHCF are collimated using a CaF_2_ lens and directed into the system. Two linear variable filters (LVFs; Vortex Optical Coatings, models 0.9–1.7–3.5–15– 0.5–2% and 1.2–2.5–3.5–15–0.5) with a filter bandwidth of 2% are used to select the desired excitation wavelengths before the beam enters the OAM system. An objective lens (0.55 NA, 40×, Olympus) focuses the filtered beam onto the sample, where optoacoustic signals are generated.

The sample is mounted on a glass coverslip, and a water tank (μ-Dish, Ibidi) with a flat glass bottom of 200 µm thickness is placed tightly above it and filled with distilled water. This setup ensures sample surface flatness and provides acoustic coupling. The combined sample and water tank assembly is held on a high-resolution XY scanning stage (8MTF-75LS05, Standa) controlled by a stepper and DC motor controller (8SMC5-USB, Standa), allowing maximum scanning speed up to 10 mm/s. A focused, immersion-type ultrasound transducer (UT, Precision Acoustics) with a central frequency of 20 MHz, 8 mm focal length, and a 10 mm aperture is submerged in the water to detect the generated acoustic signals. The transducer is coaxially aligned with the optical axis of the objective lens. The detected optoacoustic signals are filtered through analog high-pass (1 MHz) and low-pass (47 MHz) filters (Mini-Circuits), amplified by approximately 50 dB using a wideband amplifier (Spectrum Instrumentation), and collected with a high-speed digitizer (M4i.4421-x8, Spectrum Instrumentation). The digitizer operates at a 250 MS/s sampling rate with 16-bit resolution and is integrated into a control computer for data acquisition and processing.

To synchronize scanning and data collection, an external trigger delay controller (AeroDiode) is used. A small portion of the pump beam is monitored by a NIR photodetector (DET08C/M, Thorlabs), which generates a trigger signal that is fed into the delay controller. Upon receiving this signal, the controller outputs a square trigger pulse to synchronize the XY stage movement and OA signal acquisition. The data are recorded approximately 5 µs after the laser pulse and ∼0.2 µs after the square trigger signal, at a repetition rate of 1 kHz. To enhance the signal-to-noise ratio (SNR), 100 A-lines are averaged per scan point, resulting in a dwell time of approximately 83 ms.

## 3. Characterization of the system

### 3.1 Characterization of Raman fiber laser

Figure 2 (a) shows the Raman Stokes spectra generated by two pump lasers (1060 nm, blue; 1044 nm, red) with circularly polarized input light. The spectra were recorded using two optical spectrum analyzers, Yokogawa AQ6375 (0.05 nm resolution) and AQ6317B (0.015 nm resolution). Each pump wavelength produced up to eight orders of Raman Stokes lines. For the 1060 nm pump, based on the rotational Raman shift of H_2_ at 587 cm^−1^, six-orders of rotational Stokes (RS) lines were observed, spanning from ∼1100 nm to ∼1700 nm. At higher gas pressures, a vibrational Stokes (VS) line at 1894 nm emerged due to the reduced threshold of vibrational Raman gain. This was followed by a cascaded RS line generation at ∼2130 nm originating from the VS process. A similar Raman generation behavior was observed for the 1044 nm pump. Due to the circular polarization of the pump beam, RS lines exhibited higher intensities than the VS lines. Weak anti-Stokes (AS) lines extending down to ∼328 nm were also observed; however, their intensities were negligible due to the high transmission losses of the ARHCF in the UV region [32]. The discrete spectral lines generated by our system span a broad wavelength range and coincide with key chromophore absorption bands (highlighted as grey regions in Fig. 2(a)), including melanin (1110 nm), lipids (1210 nm and 1700 nm), collagen (1540 nm and 1700 nm), and water (1894 nm) [17, 33, 34]. Thanks to the narrow-linewidth pump source and the selective Raman gain profile of H_2_, each generated Raman line maintains a linewidth below 0.1 nm. This is demonstrated in Fig. 2(b) by the 2^nd^- and 3^rd^-order RS lines at ∼1278 nm and ∼1382 nm as an example, respectively. This narrow linewidth ensures high spectral purity and power density, which are essential for OA microscopy. Furthermore, the wavelength of each pulse can be finely tuned by thermally adjusting the seed laser (see Supplementary Note 2).

**Fig. 2.**
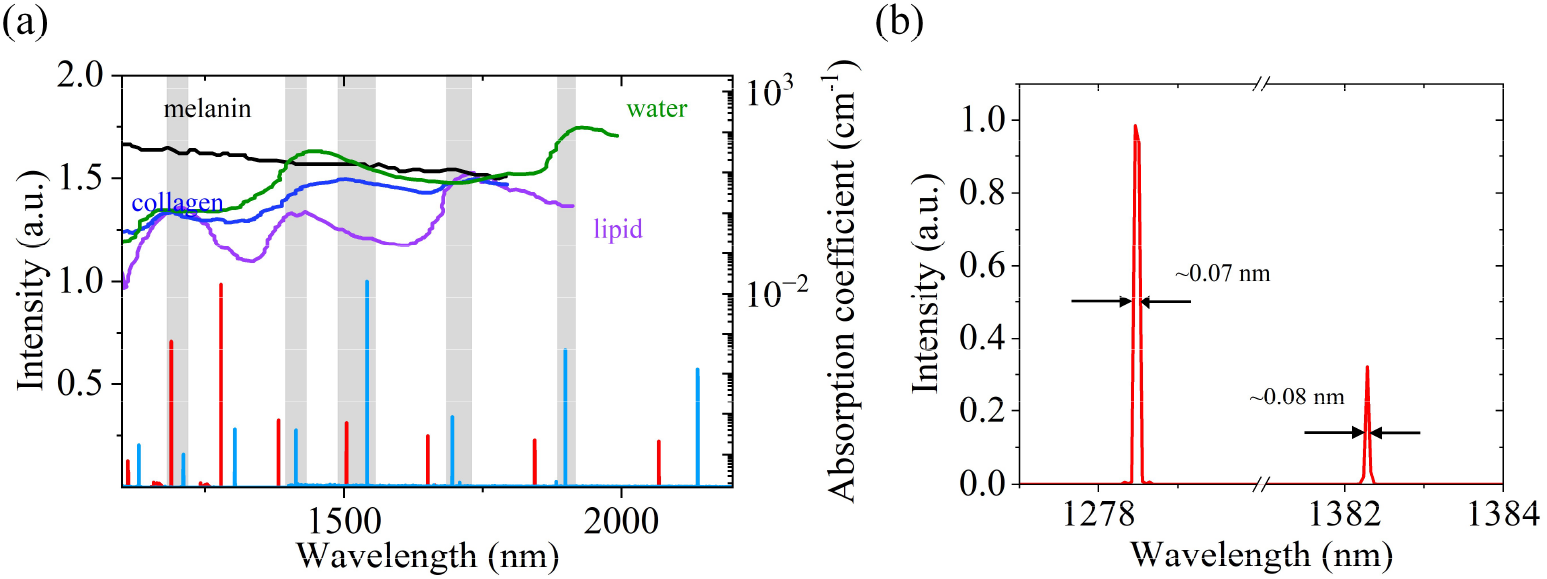
(a) Spectra of the multiwavelength Raman pulses generated using 1060 nm (blue) and 1044 nm (red) pump lasers. The right vertical axis shows the absorption spectra of water and lipids (from Ref. [33]), melanin (from Ref. [17]), and collagen (from Ref. [34]) for comparison. (b) Zoomed-in spectra of the 2^nd^-and 3^rd^-order rotational Stokes lines at ∼1278 nm and ∼1382 nm, respectively, generated by the 1044 nm pump.

OA imaging requires laser pulses with sufficient pulse energy, nanosecond scale duration, and stable output characteristics. We investigated these parameters to confirm the suitability of our laser source for OA imaging. Figures 3(a) and 3(b) show the evolution of Raman pulse energy as a function of H_2_ gas pressure. At lower pressures, only lower-order RS lines were observed. As the pressure increased, energy was progressively transferred to higher-order RS lines. At sufficiently high pressures, a vibrational Stokes (VS) line appeared, followed by additional VS-RS lines with further pressure increases. Most Raman Stokes lines exhibited pulse energies exceeding 1 μJ, except for those at 2065 nm and 2132 nm. These energy levels are sufficient for optoacoustic microscopy applications. The peak pulse energy for each Stokes line can be optimized by adjusting the gas pressure, with the optimal conditions summarized in Supplementary Note 3. For the OA imaging experiments, gas pressures between 2 and 4 bar were selected to ensure that nearly all Raman pulses maintained sufficiently high energy levels.

**Fig. 3.**
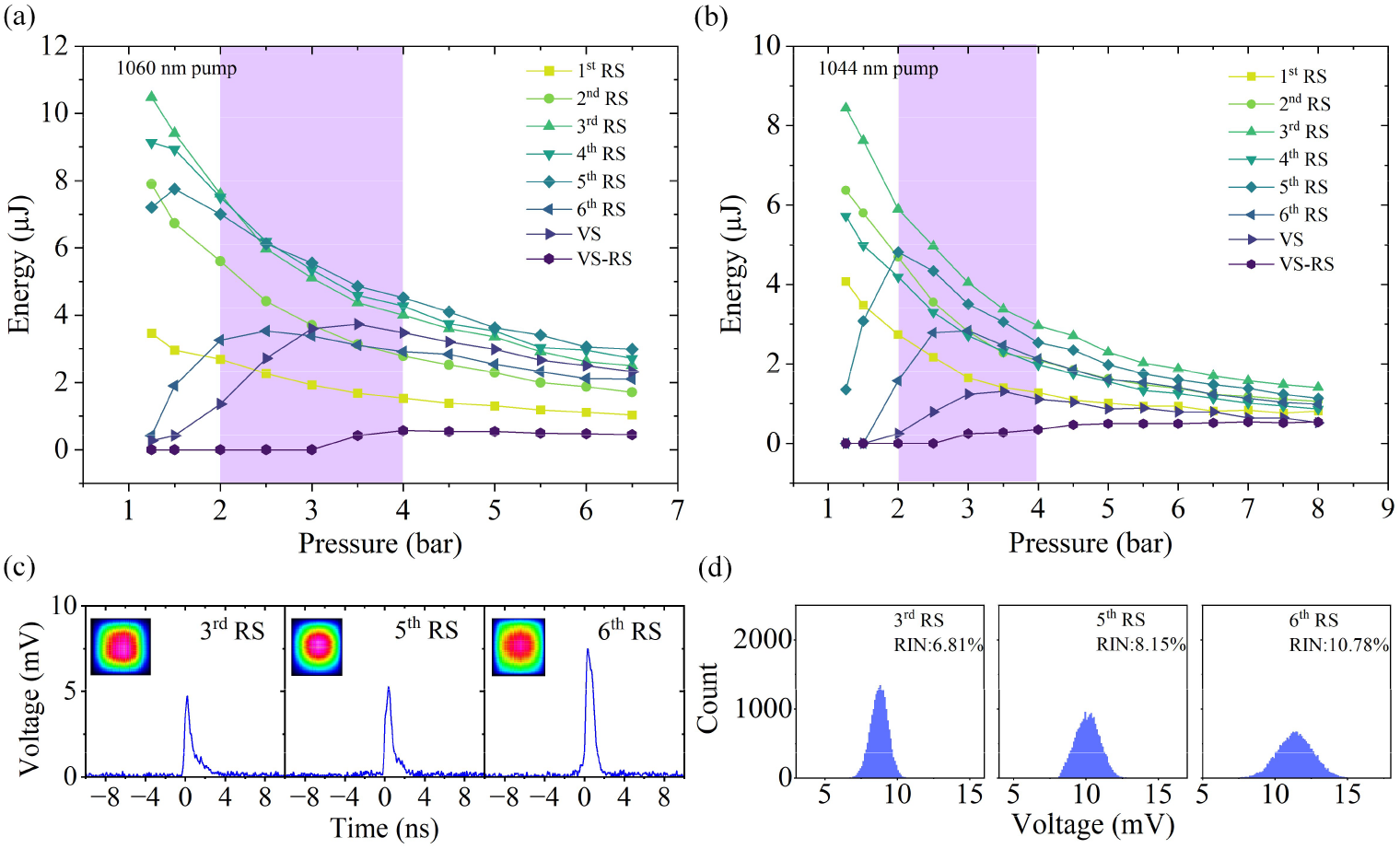
(a) Evolution of laser pulse energy for the generated Raman lines as a function of hydrogen (H_2_) pressure, using a 1060 nm pump laser. (b) Evolution of laser pulse energy for the generated Raman lines as a function of H_2_ pressure, using a 1044 nm pump laser. (c) Temporal profiles of the Raman pulses generated with the 1060 nm pump. Inset: corresponding spatial beam profiles. (d) Histograms of the pulse peak intensities for the Raman pulses generated using the 1060 nm pump.

The pulse duration and stability of the laser source were characterized, as shown in Fig. 3(c) and 3(d), using pulses generated from the 1060 nm pump as a representative example. Similar performance was observed for the 1044 nm pump. Pulse profiles were recorded at a H_2_ pressure of approximately 2 bars using a 5 GHz NIR photodetector (DET08C/M, Thorlabs) connected to a 6 GHz oscilloscope (Tektronix). The temporal profiles exhibited order-dependent characteristics, where higher-order Raman Stokes pulses displayed longer durations than lower-order ones. For instance, as shown in Fig. 3(c), the 6^th^ order RS pulse had a duration of ∼0.88 ns, compared to ∼0.76 ns for the 5^th^ order and ∼0.72 ns for the 3^rd^ order. This behavior arises because lower-order Stokes pulses act as pump sources for higher-order generation and tend to undergo temporal compression, whereas higher-order pulses are less depleted and thus maintain longer durations [35,36]. Despite these variations, all Raman pulses exhibit nanosecond-scale durations, which are optimal for efficient acoustic wave generation in OAM applications [2]. The beam quality was evaluated from the spatial profiles shown in the insets of Fig. 3(c). The Gaussian-like beam shapes indicate the laser can be focused to near-diffraction-limited spot sizes, which is critical for achieving high spatial resolution in OAM. Pulse-to-pulse stability is also a key parameter in OAM, especially for minimizing the need for signal averaging during imaging. Noise performance was quantified by calculating the relative intensity noise (RIN) from the peak intensities of 20,000 pulses at each Raman wavelength, recorded under identical pump energy conditions at 2 bar gas pressure [37]. Figure 3(d) displays representative RIN histograms for three selected Stokes orders. The RIN values measured for the 1^st^ through 6^th^ RS orders were 11.75%, 5.10%, 6.81%, 7.86%, 8.15%, and 10.78%, respectively (see Supplementary Note 4).

### 3.2 Characterization of the NIR-OAM

The lateral and axial resolutions of the NIR-OAM system were evaluated, as illustrated in Fig. 4(a) and 4(b). Lateral resolution was measured by imaging the sharp edge of a pattern on a 1951 USAF resolution target using a step size of 0.31 μm. The edge spread function (ESF) was obtained by averaging the raw OA intensity data, and the line spread function (LSF) was then derived by differentiating the ESF and applying a fitting function, as shown in Fig. 4(a). The lateral resolution, calculated from the full width at half maximum (FWHM) of the LSF, exhibited wavelength dependence consistent with theoretical predictions. Using a high-numerical-aperture objective lens, we achieved lateral resolutions ranging from approximately 1 to 2 µm, approaching the diffraction limit. Axial resolution was determined from a single OA signal obtained from the resolution target image, as shown in Fig. 4(b). The FWHM of a Gaussian fit to the Hilbert transform (HT) envelope of the signal was measured to be 30 ns, corresponding to an axial resolution of 46.2 µm, assuming a speed of sound in water of 1540 m/s.

**Fig. 4.**
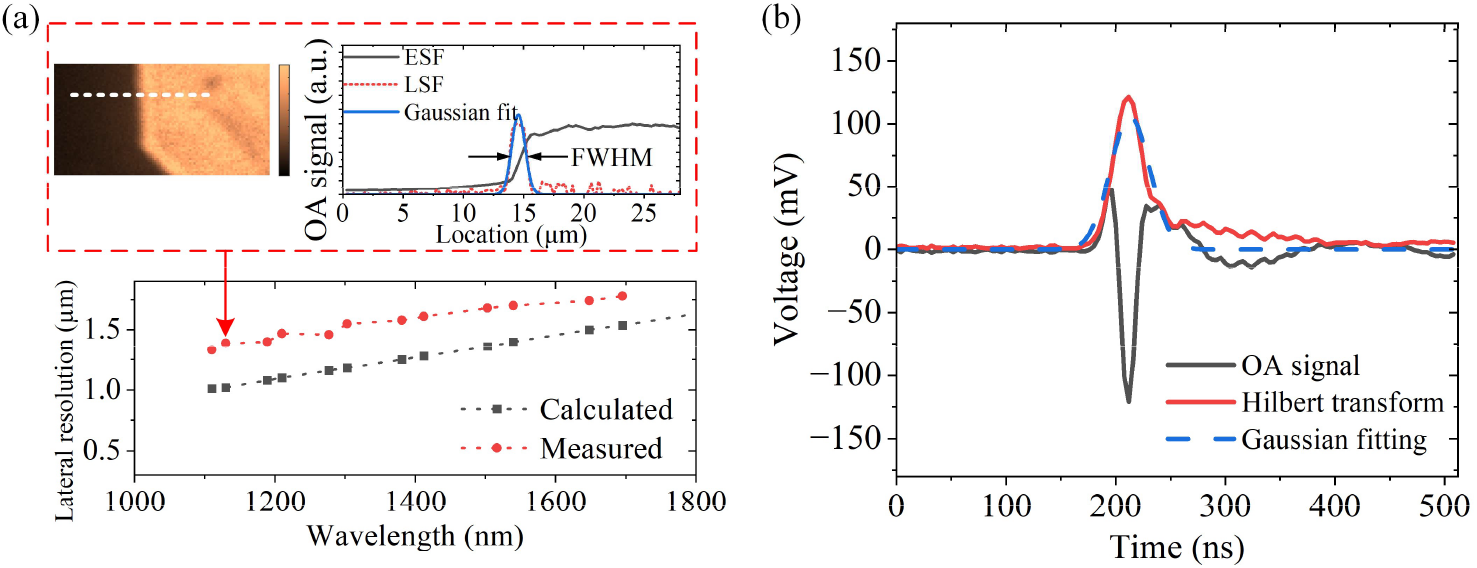
(a) Evaluated and theoretical lateral resolution of the NIR-OAM system as a function of excitation wavelength. Inset: optoacoustic (OA) image of the resolution target and corresponding Gaussian fits used to extract lateral resolution from the edge spread function (ESF) and line spread function (LSF). (b) Representative single OA signal acquired from the resolution target, along with the Hilbert transform (HT) envelope used to evaluate axial resolution.

## 4. Multi-spectral OAM

### 4.1 Mapping melanin and water

To assess the performance of the developed laser source, we first conducted spectroscopic imaging on a melanin-rich hair sample, which provides absorption across the full NIR spectrum, using surrounding water as a reference. The hair sample was placed between the water tank and coverslip, secured at both ends with tape. Deionized water was spread around the hair for acoustic coupling. Figure 5(a) shows images of the hair at various wavelengths, corrected for pulse energy variations. The imaging was performed at an 80 × 150 pixels size across a field of view (FOV) of around 200 × 375 μm^2^, corresponding to a scanning step size of 2.5 μm. For each pixel, 100 OA signals were averaged, with the absolute amplitude extracted using HT. Each scan required approximately 26 minutes. The resulting OAM images confirm wavelength-dependent absorption from the melanin in the sample, showing distinct contrasts across the broad wavelength range. The water background also demonstrates wavelength-dependent absorption, showing positive contrast around 1400-1500 nm and towards longer wavelengths.

**Fig. 5.**
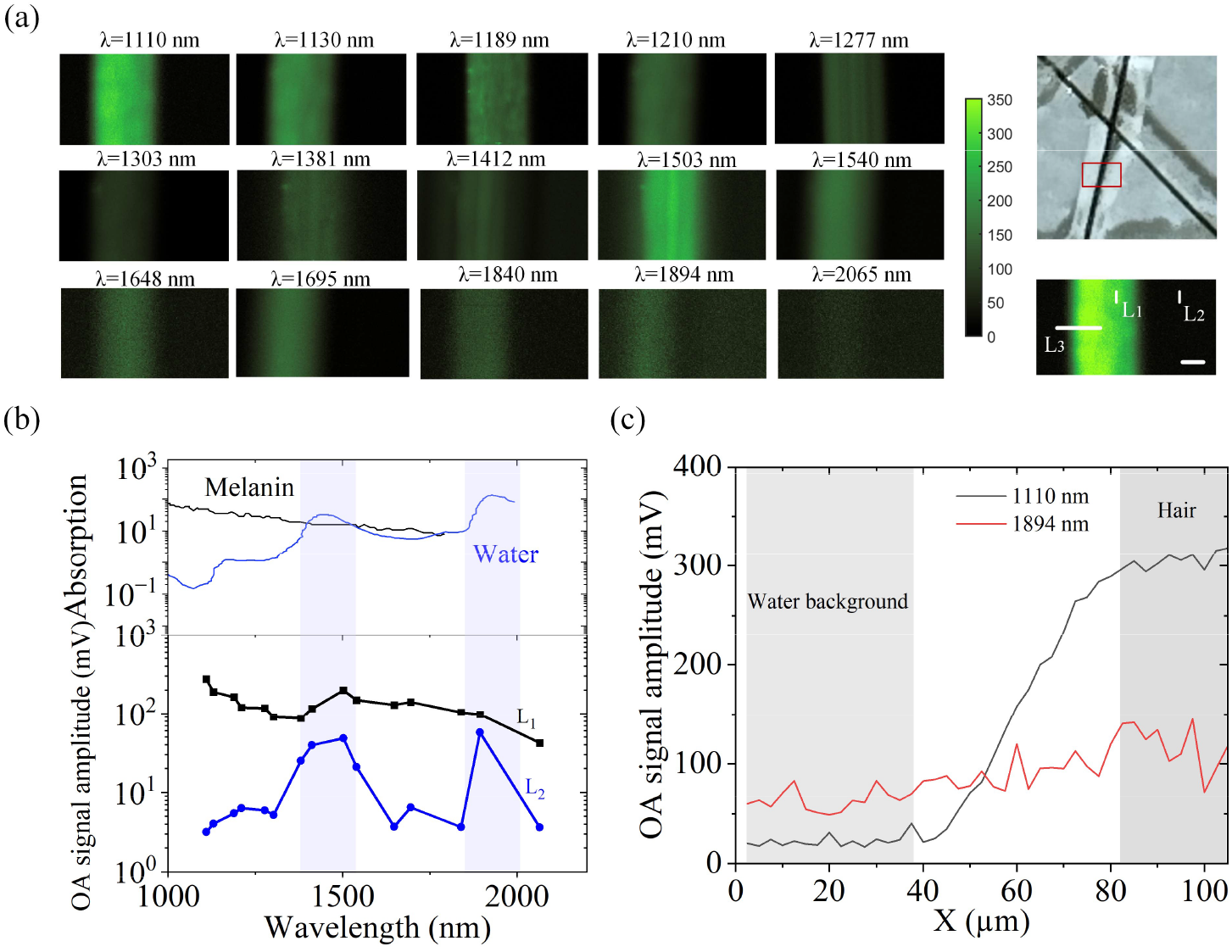
(a) OA image of hair at different wavelengths. Top figure on the right: The optical image of hair. Bottom figure in the right: OA image obtained with 1110 nm wavelength. Scale bar: 200 μm. (b) Top: Absorption spectra of melanin and water obtained from Ref. [17] and [32], respectively. Bottom: The averaged OA signal amplitudes along Line L_1_ and L_2_ as shown in (a) at different wavelengths. (c) The OA signal amplitude profile along Line L_3_ as shown in (a) at 1110 nm and 1894 nm.

To evaluate the specific absorption contrast between melanin and water, we averaged the OA signal amplitudes along line L_1_ (within the hair region) and line L_2_ (within the water region) at different wavelengths (Fig. 5(a), bottom right). The results are shown in Fig. 5(b). The overall trend in OA amplitude for melanin aligns with its corresponding absorption spectrum, with a decrease at longer wavelengths. The optimal imaging window of melanin is below 1400 nm wavelength, where water absorption is less dominant. The OA amplitudes rise from 1412 nm onward, partly due to the contribution of water absorption, which is reflected in the water background’s OA amplitude. The OA amplitude of water is maximized in two NIR bands (the light blue shaded regions), around 1400-1500 nm and c, which is consistent with water’s known absorption spectrum shown in Fig. 5(b) (from Refs. [17,32]). Additionally, the OA amplitude along line L_3_ in Fig. 5(a) at 1100 nm and 1894 nm is shown in Fig. 5(c), reflecting the large difference in absorption contrast between water and melanin at these two wavelengths.

### 4.2 Mapping gelatin patch and water

To further demonstrate the capability of the system for mapping the spatial distribution of collagen, we examined a gelatin patch as a case example, which is a hydrolyzed form of collagen and share the similar absorption peaks with collagen [34]. The imaging was conducted with a 100 × 110 pixels grid over a FOV of 2.5 × 2.75 mm^2^, with a scanning step size of 25 μm. Selecting the appropriate wavelengths is essential to effectively distinguish water and gelatin. Based on the data provided in previous literature (see Supplementary Note 5), collagen (gelatin) exhibits absorption peaks at 1200 nm, 1540 nm, 1690 nm, and 1730 nm due to CH_3_ chemical bond. Additionally, as discussed in Section 4.2, the 1400–1500 nm and 1800-2200 nm windows are optimal for imaging water content. As a comparison, the gelatin patch was imaged at two distinct wavelengths of 1412 nm and 1695 nm, as shown in Fig. 6(a). At 1695 nm, gelatin distribution is clearly visible, while the water background shows stronger intensity at 1412 nm than at 1695 nm due to the higher absorption coefficient. A composite image was also produced to facilitate the comparison, as shown in Fig. 6(a). OA signal amplitudes were quantitatively analyzed in two ROIs of 125 × 125 μm^2^ (ROI_1_ and ROI_2_ in Fig. 6(a)), as shown in Fig. 6(b). The water-dominated region (ROI_1_) exhibited higher OA amplitudes at 1412 nm, while the collagen-dominated region (ROI_2_) displayed higher amplitudes at 1695 nm, illustrating the system’s ability to map gelatin and water distributions effectively.

**Fig. 6.**
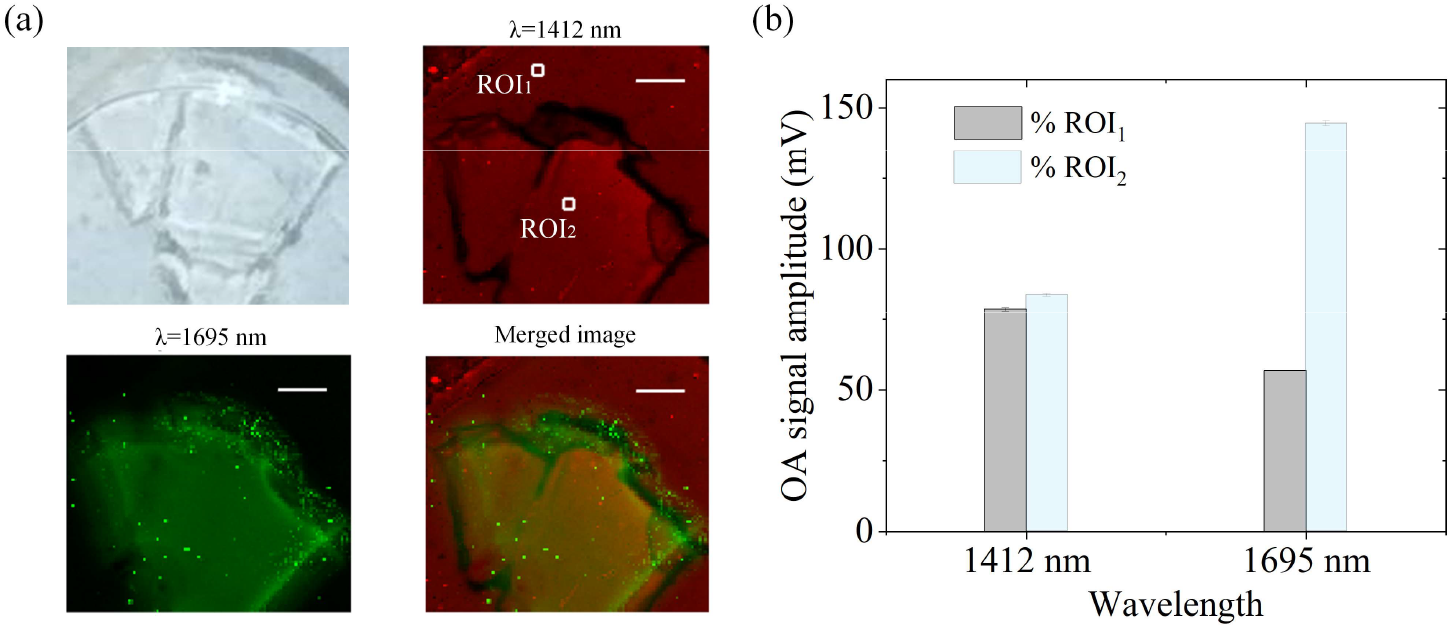
(a) Optical, OA images, and a merged OA image of a collagen patch and water surrounding. Scale bar: 500 μm. (b) The averaged OA signal amplitudes of ROI_1_ and ROI_2_ as shown in (a) at 1412 nm and 1695 nm.

### 4.3 Mapping collagen and lipid in pig tissue by spectral unmixing

To demonstrate our system’s capability to distinguish lipid and collagen, multi-spectral imaging was performed on pig skin tissue. The OA images of a 6.5 × 2.5 mm^2^ region (Fig. 7(a)) were acquired across six wavelengths with a scanning step size of 25 μm (Fig. 7(b)). Given that these two absorbers have overlapping absorption peaks due to the similar C-H bonds and simultaneously contribute to the overall OA amplitude, a spectral unmixing algorithm based on a constrained least-squares method (LLS) was employed to resolve and quantify the distribution of collagen and lipids [38]. The resulting image (Fig. 7(c)) shows lipids to be primarily distributed near the skin’s surface, while collagen was concentrated around 4 mm beneath. This result is also consistent with prior optoacoustic imaging results on pig skin tissue in Ref. [39]. The unmixing algorithm had previously validated by conducting a preliminary experiment on a hair sample at five different wavelengths and using the algorithm to distinguish the hair and water channels using our system, as shown in Fig. 7(d). The obtained OA images of the two channels and the merged OA images show that the hair signal could be clearly distinguished from the water background, confirming the suitability of the algorithm. The concordance of lipid and collagen distributions with literature and the robust phantom validation confirm the efficiency of the laser source in optoacoustic imaging of multiple biomolecules.

**Fig. 7.**
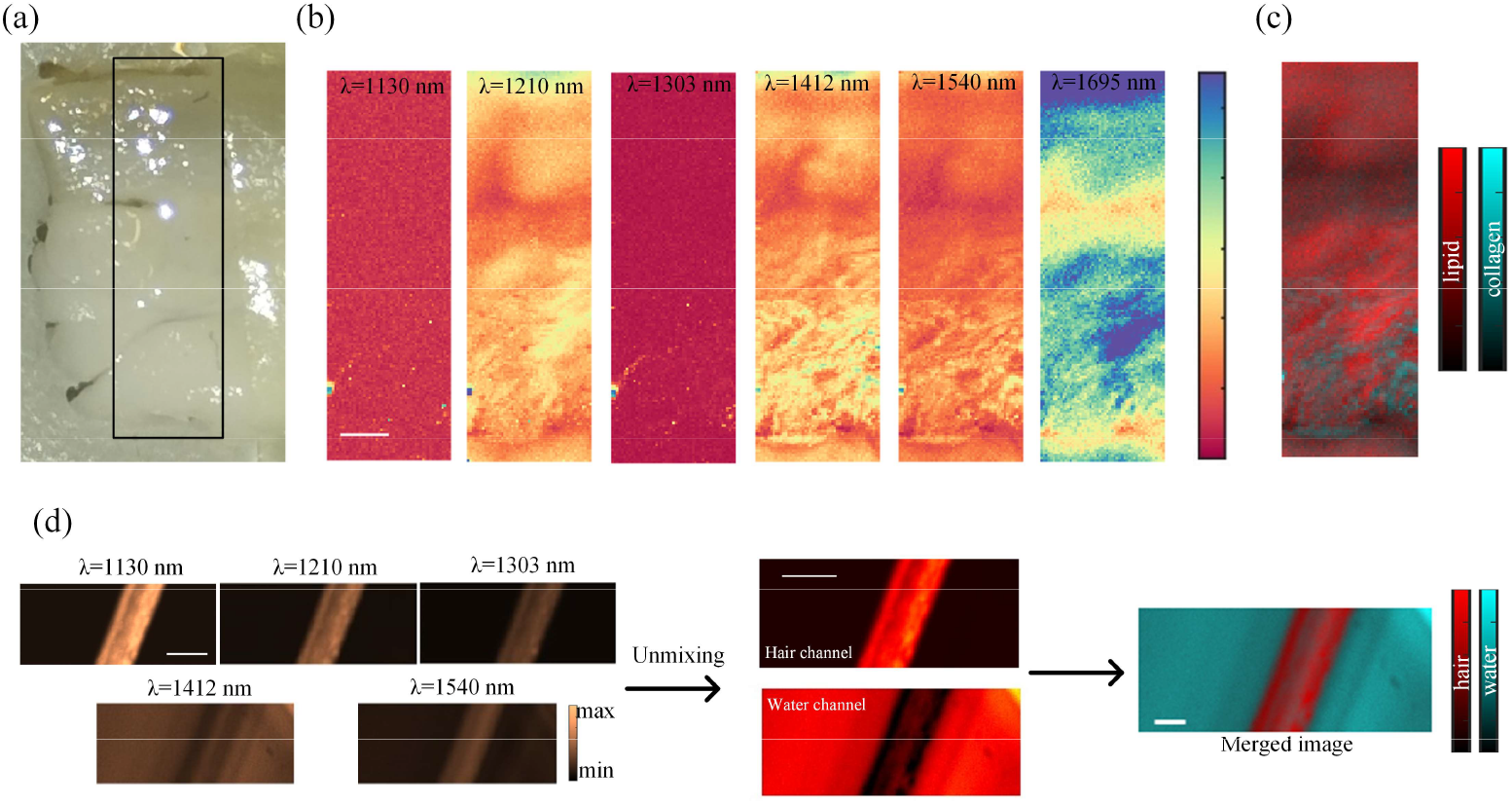
(a) Optical and (b) OA images of pig tissues. Scale bar: 500 μm. (c) Merged OA image by spectral unmixing algorithm [38]. (d) (Left) OA imaging of hair at different wavelengths. Scale bar: 100 μm. (Middle) The distinguished hair and water channels. Scale bar: 100 μm. (Right) The merged image obtained via a spectral unmixing algorithm [38]. Scale bar: 50 μm.

## 5. Conclusion

In conclusion, we have demonstrated a novel gas-based laser for multi-spectral OAM, operating across a broad spectral range from 1100 to 2200 nm. The system is based on the emerging gas-filled ARHCF platform. Each Raman laser line delivers μJ-level pulse energy, nanosecond-scale pulse durations, and a narrow spectral linewidth of less than 0.1 nm. The wavelengths can be selectively tuned to overlap with the characteristic absorption peaks of key biological chromophores, including lipids (1210 nm and 1700 nm), collagen (∼1700 nm), and water (∼1400 nm and 1900 nm). Consequently, the spatial distributions of these chromophores within biological samples can be effectively resolved using our system. Overall, the developed multiwavelength fiber laser represents a promising and versatile light source for advanced OAM applications.

## Data availability

Data underlying the results presented in this paper are available upon request.

## Supplemental document

See Supplementary Document for supporting content.

