## Supplementary material for "Multi-spectral optoacoustic microscopy driven by gas-filled hollow-core fiber laser pulses": Supplymentary Material

#### **Supplementary Note 1.** Review of the multispectral fiber laser sources used for near-infrared (NIR) optoacoustic microscopy (OAM).

**Table S1. Review of the multispectral fiber laser sources used for NIR-OAM.**

| Ref | Laser source | Wavelength (nm) | Target Chromophore | Energy (μJ) | Linewidth (nm) | Resolution (μm) |
|---|---|---|---|---|---|---|
| [1] | SC laser | 1440-1870 | Lipid | 0.197 | 33-38 | ~11.46 μm (1720 nm) |
| [2] | | 1650-1850 | Lipid | 0.2~0.6 | 38-42 | ~11.8 μm |
| [3] | | 475-2400 | Hb, melanin, lipid, collagen, glucose, | 0.017–0.11 | 25 | 5-7 μm |
| [4] | SRS-based fiber laser | 1064-1325 | Lipid | 0.20-1.17 | 4-55 | 6.2 μm |
| [5] | | 1050-1325 | Lipid | 1~6 (0.32 for 1720) | 10-50 | / |
| [6] | | 1098-1270 | Lipid | 3~60 | 10-50 | 43-63 μm |
| [7] | | 1168.4, 1202.1 | polymer | >3 μJ | 0.07 | / |
| [8] | | 1700.2, 1710.4, 1720.3 | polymer | 1.1 | 0.173 | / |
| This work | Gas-filled ARHCF laser | 1110-2165 | Melanin, lipid, collagen | >1 for each pulse | <0.1 | 1-2 μm |

[a]PRR: Pulse repetition rate; SC: super continuum; SRS: Stimulated Raman Stokes; ARHCF: Anti-resonant hollow-core fiber.

#### **Supplementary Note 2.** Wavelength tunability range of the Raman lines

**Table S2. Calculated wavelength range of the Raman lines.**

| | Seed 1: (1044-1045 nm) | Seed 2: (1060-1061 nm) |
|---|---|---|
| $1^{st}$ -order RS | 1112.2-1113.3 | 1130.3-1131.5 |
| $2^{nd}$ -order RS | 1189.9-1191.1 | 1210.6-1212 |
| $3^{rd}$ -order RS | 1279.3-1280.6 | 1303.2-1304.8 |
| $4^{th}$ -order RS | 1383.2-1384.7 | 1411.1-1413 |
| $5^{th}$ -order RS | 1505.4-1507.2 | 1538.5-1540.8 |
| $6^{th}$ -order RS | 1651.3-1653.5 | 1691.2-1694 |
| VS | 1843.8-1846.9 | 1894.3-1897.5 |
| $1^{st}$ -order RS-VS | 2053.2-2057 | 2116-2120 |

### Supplementary Note 3. Energy characterization of the Raman pulses

**Table S3. Maximum energy of Raman pulses and corresponding gas pressure.**

| Pump laser | Wavelength (nm) | Maximum energy (μJ) | Throughput to coupled pump (%) | Pressure (bar) |
|---|---|---|---|---|
| 1044 nm | 1110 | 4.07 | 8.75 | 1.25 |
| | 1189 | 6.36 | 13.67 | 1.25 |
| | 1277 | 8.44 | 18.15 | 1.25 |
| | 1381 | 5.72 | 12.3 | 1.25 |
| | 1503 | 4.81 | 10.34 | 2 |
| | 1648 | 2.83 | 6.08 | 2.5 |
| | 1840 | 1.308 | 2.81 | 3.5 |
| | 2065 | 0.54 | 1.16 | 7 |
| 1060 nm | 1130 | 3.45 | 5.67 | 1.25 |
| | 1210 | 7.89 | 12.98 | 1.25 |
| | 1303 | 10.48 | 17.24 | 1.25 |
| | 1410 | 9.13 | 15.02 | 1.25 |
| | 1540 | 7.74 | 12.73 | 1.5 |
| | 1695 | 3.52 | 5.79 | 2.5 |
| | 1876 | 3.72 | 6.12 | 3.5 |
| | 2132 | 0.56 | 0.92 | 4 |

#### Supplementary Note 4. Pulse profiles and Relative intensity noises (RIN) of the Raman pulses

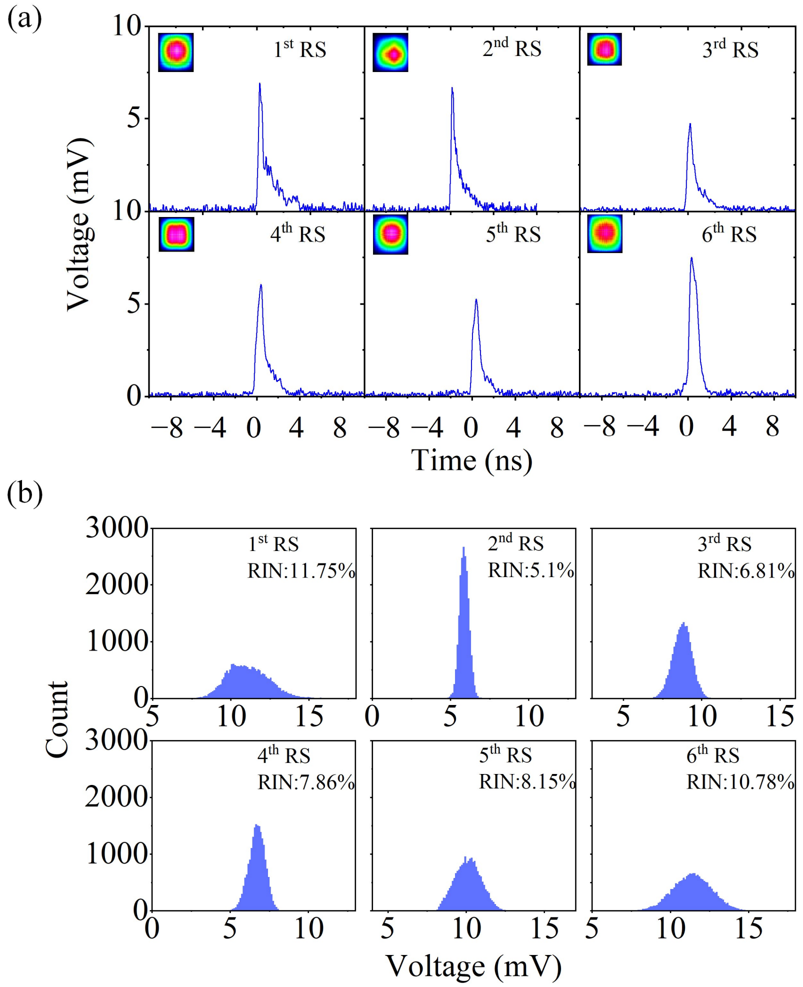


Fig. S1 (a) Pulse profiles and (b) Relative intensity noises (RINs) of the $1^{st}$ to $6^{th}$-orders Raman pulses.

### Supplementary Note 5. Absorption peaks of collagen from references

**Table S4. Summary of absorption peak of collagen.**

| Ref | Absorption peak wavelength (nm) |
|---|---|
| [9] | 1700-1730 |
| [10] | 1200 |
| [11] | 1720 |
| [12] | 1200, 1500 |
| [13] | 1600 |
| [14] | 1200, 1550, 1700 |
| [15] | 1200 1550 1700 |
